# *MYBPC3* Mutations cause Hypertrophic Cardiomyopathy by Dysregulating Myosin: Implications for Therapy

**DOI:** 10.1101/304204

**Authors:** Christopher N. Toepfer, Hiroko Wakimoto, Amanda C. Garfinkel, Barbara McDonough, Dan Liao, Jianming Jiang, Angela Tai, Josh Gorham, Ida G. Lunde, Mingyue Lun, Thomas L. Lynch, Sakthivel Sadayappan, Charles S. Redwood, Hugh Watkins, Jonathan Seidman, Christine Seidman

**Affiliations:** Department of Genetics, Harvard Medical School, Boston, MA, USA.; Division of Cardiovascular Medicine, Radcliffe Department of Medicine, University of Oxford; Wellcome Centre for Human Genetics, University of Oxford.; Cardiology, Children’s Hospital Boston, Boston, MA, USA.; Department of Biochemistry & Cardiovascular Research Institute(CVRI), Yong Loo Lin School of Medicine, National University of Singapore, Singapore.; Institute for Experimental Medical Research, Oslo University Hospital and University of Oslo, Oslo, Norway.; Department of Medicine, Division of Genetics, Brigham and Women’s Hospital, Boston, Massachusetts, USA.; Department of Molecular Pharmacology and Therapeutics, Health Sciences Division, Loyola University Chicago, Maywood, Illinois 60153; Division of Cardiovascular Health and Disease, Heart, Lung and Vascular Institute, Department of Internal Medicine, University of Cincinnati, Cincinnati, OH, USA; Howard Hughes Medical Institute, Brigham and Women’s Hospital, Boston, MA, USA

## Abstract

The mechanisms by which truncating mutations in *MYBPC3* (encoding cardiac myosin binding protein-C; cMyBPC) or myosin missense mutations cause hyper-contractility and poor relaxation in hypertrophic cardiomyopathy (HCM) are incompletely understood. Using genetic and biochemical approaches we explored how depletion of cMyBPC altered sarcomere function. We demonstrate that stepwise loss of cMyBPC resulted in reciprocal augmentation of myosin contractility. Direct attenuation of myosin function, via a damaging missense variant (F764L) that causes dilated cardiomyopathy (DCM) normalized the increased contractility from cMyBPC depletion. Depletion of cMyBPC also altered dynamic myosin conformations during relaxation - enhancing the myosin state that enables ATP hydrolysis and thin filament interactions while reducing the super relaxed conformation associated with energy conservation. MYK-461, a pharmacologic inhibitor of myosin ATPase, rescued relaxation deficits and restored normal contractility in mouse and human cardiomyocytes with *MYBPC3* mutations. These data define dosage-dependent effects of cMyBPC on myosin that occur across all phases of the cardiac cycle as the pathophysiologic mechanisms by which *MYBPC3* truncations cause HCM. Therapeutic strategies to attenuate cMyBPC activity may rescue depressed cardiac contractility in DCM patients, while inhibiting myosin by MYK-461 should benefit the substantial proportion of HCM patients with *MYBPC3* mutations.

**One Sentence Summary:** Analyses of cardiomyocytes with hypertrophic cardiomyopathy mutations in *MYBPC3* reveal that these directly activate myosin contraction by disrupting myosin states of relaxation, and that genetic or pharmacological manipulation of myosin therapeutically abates the effects of *MYBPC3* mutations.

## Introduction

Hypertrophic cardiomyopathy (HCM) is a heritable disease of heart muscle affecting ∼ 1 in 500^1^ individuals. Patient symptoms can be minimal or relentlessly progressive with resultant heart failure and/or sudden cardiac death^2^. Adverse clinical outcomes in HCM increase with disease duration, thereby underscoring the importance of therapeutic strategies to abate disease progression^3^.

Dominant pathogenic variants in eight sarcomere genes cause HCM, but predominate in *MYBPC3* and *MYH7* (encoding β cardiac myosin heavy chain; β-MHC) ^4^. The overwhelming majority of HCM founder mutations^5–11^, including one affecting 4% of South Asians^12^ reside in *MYBPC3*. All HCM mutations in *MYH7* encode missense substitutions^4^ and mutant myosins are incorporated into the sarcomere. By contrast, most *MYBPC3* mutations are truncating and are predicted to cause haploinsufficiency of cMyBPC^13, 14^ The mechanisms by which distinctive mutations in these two sarcomere proteins uniformly produce hyperdynamic contraction and poor relaxation (diastolic dysfunction) in advance of the morphologic remodelling in HCM^15–17^ remain incompletely understood ^18^.

Biophysical analyses demonstrate that HCM mutations in β-MHC, the molecular motor of the sarcomere, increase ATPase activity, actin-sliding velocity, and power (reviewed^19^). Structural data also predict that *MYH7* mutations disrupt dynamic conformational interactions that occur between paired myosin molecules during relaxation^20^. These are denoted as i) disordered relaxation (DRX), a state where one or both myosin heads could be active, able to hydrolyse ATP and potentiate force; and ii) super relaxation (SRX), a state of dual inactivation of myosins with both ATPases inhibited.

cMyBPC has structural and functional roles in sarcomere biology ^21^. cMyBPC is generally thought to serve as a brake that limits cross bridge interactions^21^ through its biophysical interactions of its amino and carboxyl termini with both myosin^22^ and actin^23^. Phosphorylation of the amino terminus of cMyBPC reduces myosin and increase actin interactions to promote cross-bridge formation^22^, events that are reversed by calcium levels that maximally activate thin filaments^23, 24^ Interpreting these interactions in the context of HCM mutations that reduce cMyBPC levels is complex for several reasons. Cardiac histopathology and *in vivo* function of heterozygous Mybpc3^+/-^ mice, which genetically recapitulate human HCM mutations, are indistinguishable from wildtype. Homozygous loss of cMyBPC impairs developmental cytokinesis resulting in expansion of mononuclear cardiomyocytes^25^ and cardiac function that mimics cMyBPC phosphorylation, with increased contractile force^23, 26^. Recent studies also demonstrate that loss of cMyBPC also alter proportions of relaxed myosin in DRX and SRX conformations^27, 28^, but whether or not this relates to contractility is unknown.

To better understand how *MYBPC3* mutations cause HCM, we assessed sarcomere function in the setting of cMyBPC deficiency and genetically altered myosin or pharmacologic attenuation of myosin ATPase activity. In combination, these assays shed light on unifying mechanisms that drive HCM pathophysiology and demonstrate that a single pharmacologic manipulation of myosin corrects sarcomere dysfunction caused by *MYBPC3* mutations, thereby providing a promising avenue for treating this prevalent cause of human HCM.

## Results

We studied three mouse models with altered cMyBPC expression (Supplemental Figure 1). Mybpc3^t/+^ and Mybpc3^t/t^ mice carry endogenous heterozygous or homozygous truncating mutations^29, 30^ and express graded reductions of cMyBPC protein. MyBPC-RNAi^25^ are wildtype (WT) mice transfected (P10) with a cardiotropic adeno-associated (serotype 9) virus carrying green fluorescent protein (GFP) and RNAi targeting *Mybpc3* transcripts. Injection of 5 × 10^11^ viral genomes (vg)/kg reduced *Mybpc3* transcripts to less than 10% of levels in WT mice and abolished protein expression (Supplemental Figures 1-3). Sham-RNAi +/- denotes Mybpc3^t/+^ mice transfected with virus carry GFP alone. We also studied heterozygous (Myh6^F764L/+^) and homozygous (Myh6^F764L/ F764L^) mice, which mimic a human *MYH7* mutation that causes dilated cardiomyopathy (DCM).^31, 32^

### In vivo cardiac phenotypes in Mypbc3 mutant mice

Previous studies demonstrate that Mybpc3^t/+^ mice have normal contractility and morphology, but with increased load Mybpc3^t/+^ mice develop greater left ventricular hypertrophy than WT mice ^29, 33^. Mybpc3^t/t^ mice have significantly increased LV volumes and hypertrophy (increased LV wall thickness; LVWT) due in part to increased numbers of cardiomyocytes from additional perinatal divisions prior to permanent exit from the cell cycle.^25^ By contrast, MyBPC-RNAi mice developed progressive hypertrophy in comparison WT (Supplemental Figure 2A-C) similar to other HCM mice^25^, without changes in ventricular volumes. Fractional shortening, an *in vivo* measure of contractility, was not significantly different between MyBPC-RNAi and WT mice.

### Contractility and Relaxation in Cardiomyocytes from Mypbc3 Mutant Mice

As in vivo contractility and relaxation reflects sarcomere performance as well as myocardial geometry, histopathology, and hemodynamic load, we studied *ex vivo* cardiomyocytes to assess biophysical functions of sarcomeres with altered cMyBPC levels. Isolated cardiomyocytes from at least four mice of each genotype (cells numbers indicated in figure legends) were studied. Sarcomere length was measured throughout the contractile cycle to assess cellular shortening, a surrogate systolic function (Figure 1A-D). Cardiomyocytes isolated from naïve or sham-RNAi +/- mice had comparable cell shortening thereby excluding an effect of AAV9 on contractility (Supplemental Figure 2D). Cardiomyocytes with altered cMyBPC expression had dosage-dependent increases in maximal cellular shortening (Figure 1C). In comparison to WT, Mybpc3^t/+^ cardiomyocytes had 50% increased shortening (7.2 ± 0.25% p < 0.03), whilst cellular shortening was increased by 100% in cardiomyocytes lacking cMyBPC (MyBPC-RNAi: 10 ± 0.6%; Mybpc3^t/t:^ 9.5 ± 0.9%; p<0.0001). Notably, augmentation of cellular shortening of isolated cardiomyocytes with cMyBPC deficiency did not result in increased contractility in vivo,^25, 29^ an observation that implies additional (biochemical, transcriptional, and morphologic) processes can modulate ensemble systolic performance of cardiomyocytes.

**Figure 1:**
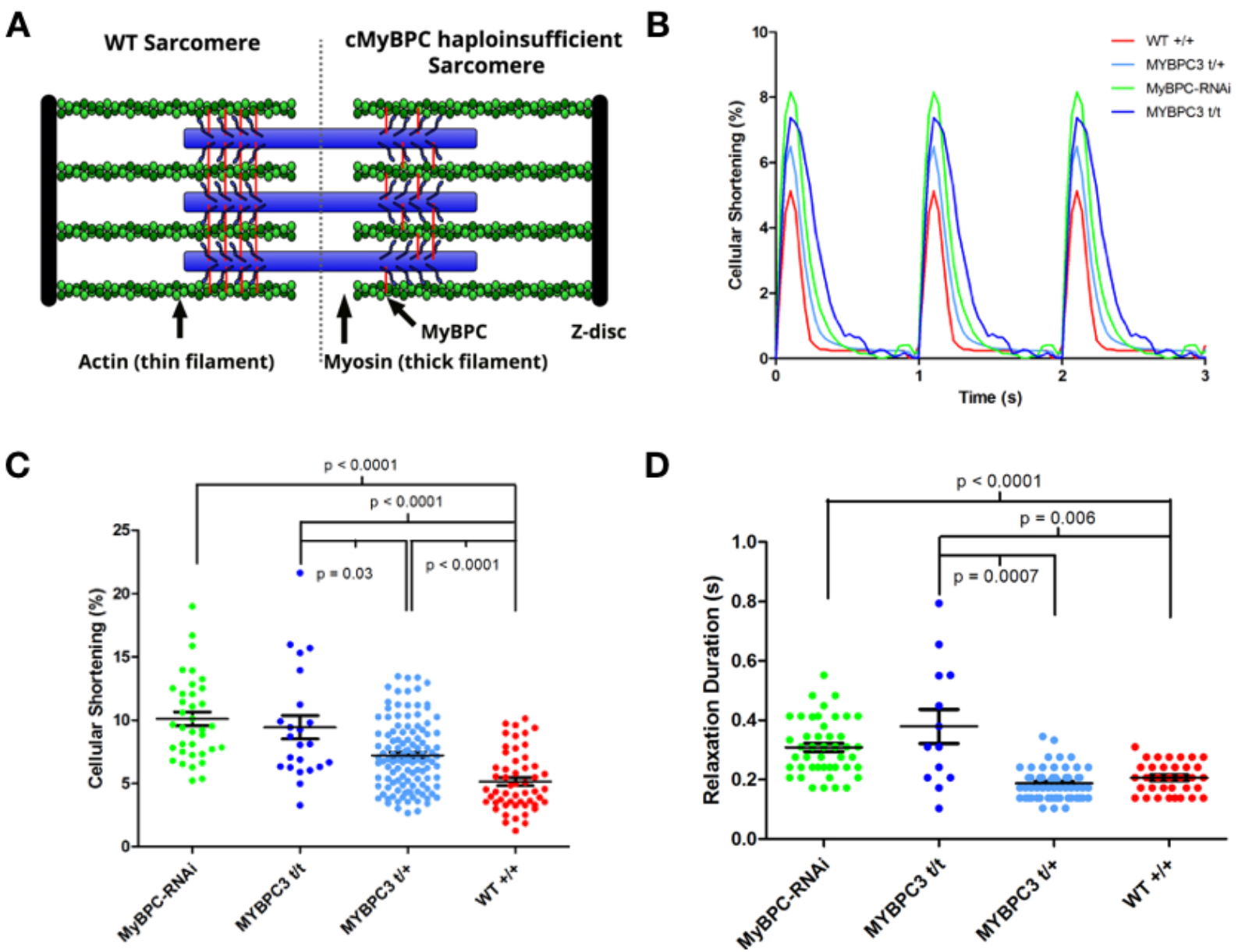
Contractile characterization of cMyBPC mouse models. **A)** A schematic depiction of the WT sarcomere with normal cMyBPC integration (left half) and the consequences of mutations that deplete cMyBPC levels in the sarcomere (right half). **B)** Representative contractile waveforms from isolated cardiomyocytes paced at 1Hz. Sarcomere lengths of isolated cardiomyocytes were tracked to define the percentage shortening per cell and duration of relaxation. Each trace is the averaged waveform across all cells analyzed for each treatment group. **C)** Comparisons of cellular shortening of isolated cardiomyocytes from four mice with different genotypes. (Cells analyzed: MyBPC-RNAi = 36; Mybpc3^t/t^ = 23; Mybpc3^t/+^ = 118,WT = 53.) Data is plot as mean ± SEM. **D)** Measures of duration from peak contraction to relaxation in seconds plot as mean ± SEM (Cells analyzed: MyBPC-RNAi = 34; Mybpc3^t/t^ = 13; Mybpc3^t/+^ = 61,WT = 30).

Human HCM is characterized by impaired diastolic performance, a parameter that is difficult to assess in mice. However, tracking sarcomere lengths in isolated cardiomyocytes across the contractile cycle provided a quantitative proxy for diastolic function, the duration of relaxation (Figure 1D). Relaxation was prolonged in MyBPC-RNAi and Mybpc3^t/t^ cardiomyocytes (0.31 ± 0.02; p < 0.0001 and 0.38 ± 0.06; p < 0.0056) compared to WT cells (0.21 ± 0.02), while the duration of relaxation in Mybpc3^t/+^ cardiomyocytes was indistinguishable from WT (0.19 ± 0.02).

### Genetic Repression of Myosin Function Corrects cMyBPC Deficiency

Rare myosin missense mutations cause DCM, a disorder characterized by ventricular enlargement and diminished cardiac contractility Mice engineered to carry the human DCM mutation Myh6^F764L/+^ and Myh6^F764L/ F764L^ mice recapitulate these phenotypes.^31^ Analyses of isolated cardiomyocytes from these models showed a genotype-dependent depressed cellular shortening (Figure 2A): contractility in Myh6^F764L/+^ and Myh6^F764L/ F764L^ cardiomyocytes was 75% and 50% of normal (P < 0.0001 for each). DCM cardiomyocytes also had small, but significantly reduced durations of relaxation (Figure 2B) (Myh6^F764L/+^: 0.17 ± 0.01, p = 0.0002; Myh6^F764L/ F764L^: 0.14 ± 0.005, p < 0.0001) compared to WT cardiomyocytes (0.21 ± 0.02).

**Figure 2:**
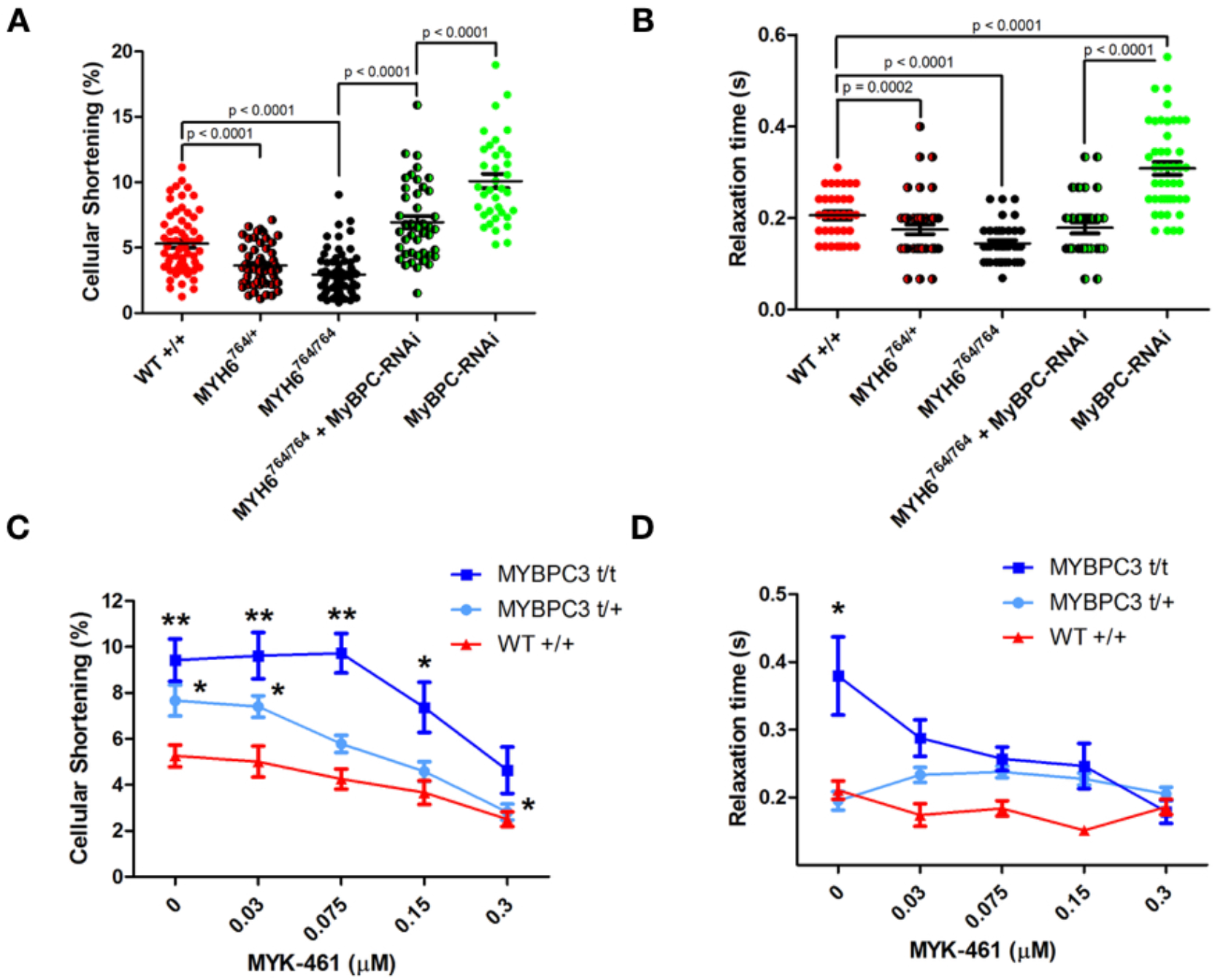
Genetic and pharmacological manipulation of cardiomyocytes depleted for cMyBPC. **A)** Sarcomere contractility in isolated cardiomyocytes from WT, Myh6^764/+^, Myh6^764/764^, Myh6^764/764^ + MyBPC-RNAi, and MyBPC-RNAi mice. (Cells analyzed; WT = 63, Myh6^764/+^ = 55, Myh6^764/764^ = 71, Myh6^764/764^ + MyBPC-RNAi = 43, MyBPC-RNAi = 36) **B)** Sarcomere relaxation of isolated cardiomyocytes from WT, Myh6^764/+^, Myh6^764/764^, Myh6^764/764^ + MyBPC-RNAi, and MyBPC-RNAi mice. Individual data points are plot with mean ± SEM indicated. All significant p values are indicated on the graph. (Cells analyzed; WT = 30, Myh6^764/+^ = 45, Myh6^764/764^ = 29, Myh6^764/764^ + MyBPC-RNAi = 31, MyBPC-RNAi = 45) **C)** Sarcomere contractility of cardiomyocytes treated with 0.03 – 0.3 μM MYK-461. More than 20 cardiomyocytes were analyzed for each drug concentration and treatment group) **D)** Sarcomere relaxation of cardiomyocytes treated with 0.03 – 0.3 μM MYK-461. All data is displayed as mean ± SEM. *p < 0.01 and **p < 0.0001 denote comparisons with WT without MYK-461.

To determine if manipulation of myosin properties would alter hyper-contractility of *Mybpc3* mutations, we injected AAV9 carrying GFP and MyBPC-RNAi into P10 Myh6^F764L/ F764L^ mice. Forty days post injection the cellular shortening (Figure 2A) and duration of relaxation (Figure 2B) were significantly improved in comparison to MyBPC-RNAi cardiomyocytes (p<0.0001) and both contractile and relaxation parameters were indistinguishable from WT cardiomyocytes.

### Inhibition of Myosin ATPase Corrects cMyBPC Defects in Cardiomyocytes and Cardiac Tissues

As genetic deficits in myosin normalized the performance of *Mybc3*-deficient cardiomyocytes, we tested the hypothesis that MYK-461, a cardiac-selective, pharmacologic allosteric myosin ATPase inhibitor, would also be effective. Acute treatment of Mypbc3^t/+^ and Mypbc3^t/t^ cardiomyocytes with MYK-461 reduced the hyper-contractility in a dose dependent manner (Figure 2C). Contractile function was normalized in Mypbc3^t/+^ cardiomyocytes at 0.15 μM MYK-461 and in Mypbc3^t/t^ dosages at ∼ 0.3 μM. The higher dose of MYK-461 reduced sarcomere contractility by ∼50% in both mutant genotypes and depressed contractility by ∼30% in WT cardiomyocytes. Concurrently, 0.3 μM of MYK-461 normalized relaxation times in Mypbc3^t/t^ cardiomyocytes (Figure 2D), but did not alter the duration of relaxation in Mypbc3^t/+^ or WT cardiomyocytes.

### Increased Ratios of Myosin Heads in DRX:SRX caused by cMyBPC-deficiency are Normalized by Modulation of Myosin ATPase

The proportions of myosin heads in DRX and SRX conformations correlate with the rate of ATP cycling in relaxed muscle, which can be measured by the decay of a fluorescent, non-hydrolyzable ATP (Mant-ATP) from skinned muscle^34^ or cell fibers. (Figure 3 and Supplemental Figure 4). Myosin heads in the DRX configuration have ∼5× more ATPase activity than myosin heads in the SRX configuration^34^. Hence the fraction of myosin heads in the SRX configuration and DRX configuration can be estimated from the fraction of Mant-ATP that is released rapidly (DRX) or slowly (SRX).

**Figure 3:**
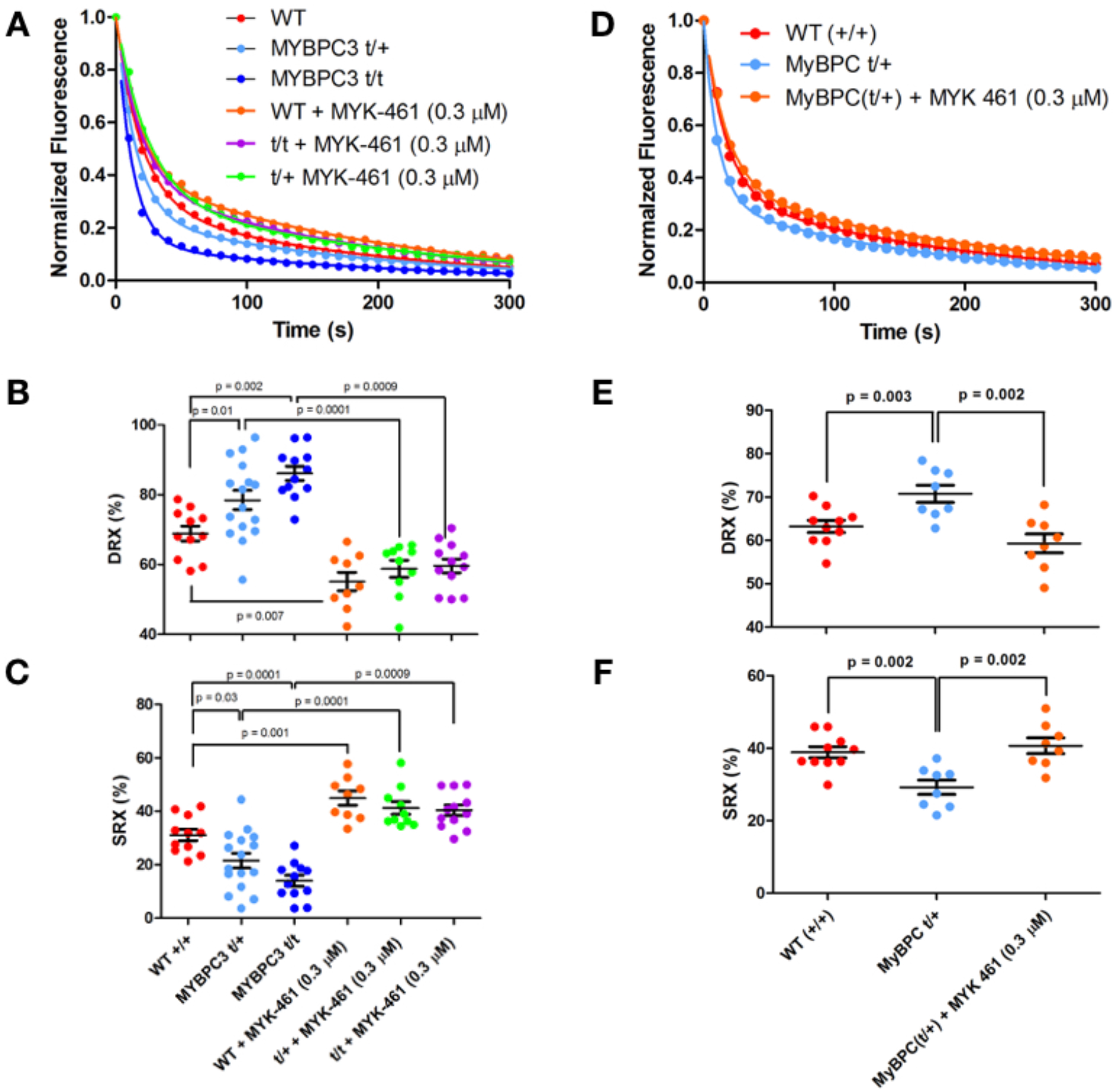
Mant-ATP assessment and correction of SRX and DRX ratios in mouse and human myocardium. **A)** Average Mant-ATP fluorescence decay curves plot from fluorescence decay due to dark ATP wash, acquisition duration 5 minutes. Data points are the mean of ∼12 separate experiments from 3 separate individuals in each genotype/treatment group. Data is fit by a double exponential decay to assess ratios of DRX and SRX heads (Methods). **B)** Plot of the initial rapid decay amplitude corresponding to DRX heads. **C)** Plot of the second exponents slow decay amplitude corresponding to SRX heads. **D)** Average Mant-ATP fluorescence decay curves of unrelated human hearts: three without HCM (WT) and three HCM heart with *MYBPC3* t/+ mutations. Each curve is the average of 12 experiments from three separate samples in each treatment group. Data is fit by double exponential decay to assess ratios of DRX and SRX heads in the myocardium. **E)** Plot of the initial rapid decay amplitude corresponding to DRX heads. **F)** Plot of the second exponents slow decay amplitude corresponding to SRX heads. All data is presented in each panel, plotted as mean ± SEM with significances indicated with p values.

Assays of skinned cardiac fibers from WT, Mypbc3^t/+^ and Mypbc3^t/t^ mice showed significantly different proportions of myosins in DRX and SRX confirmations (Figure 3 A-C). Compared to WT cardiac tissues, Mypbc3^t/+^ and Mypbc3^t/t^ had a 12% shift (p = 0.0274) and 23% shift (p = 0.0002), respectively, of myosins in DRX (Figure 3B), changes that paralleled the dose-dependent increase in cardiomyocyte contractility (Figure 1C). By contrast, assessment of the proportions of SRX and DRX found in Myh6^F764L/+^ and Myh6^F764L/F764L^ were comparable to those of wildtype mice, a finding that indicates that this DCM mutation should impact only contractility with minimal impact on cardiomyocyte relaxation (Supplemental Figure 5).

As an allosteric inhibitor of myosin ATPase, we asked if MYK-461 would also alter the fraction of myosin heads in the SRX and DRX configuration in cardiac tissues from WT and cMyBPC-deficient mice. MYK-461 (0.3 μM) treatment of skinned cardiac fibers (Figure 3 A-C) increased the proportion of myosins in SRX and reduced myosins in DRX by 15% in WT (p = 0. 007), 20% in Mypbc3^t/+^ (p = 0.001) and by 26% in Mypbc3^t/t^ fibers (p = 0.0009) in comparison to untreated corresponding genotypes. Notably this MYK-461 dose normalized cardiomyocyte contractility (Figure 2C,D), indicating a direct relationship between the proportion of myosins in DRX and cellular hyper-contractility.

Analyses of Mant-ATP release from skinned human heart fibers with heterozygous *MYBPC3* truncations showed abnormalities comparable to those in mutant mouse hearts. The proportion of myosin in DRX was increased (∼12%, p = 0.0031) compared to normal human heart fibers (Figure 3 D-F). Moreover, treatment of *MYBPC3*-deficient fibers with 0.3 μM MYK-461 normalized the ratio of DRX/SRX, by reciprocally reducing the proportion in DRX and increasing the proportion in SRX by 16% (p = 0.0022, vs. untreated).

Taken together, these observations indicate that *MYBPC3* mutations in humans and mice disrupt normal myosin conformations, resulting in the increased contractility and ATP consumption associated with HCM. These abnormalities can be pharmacologically corrected with the myosin ATPase inhibitor, MYK-461.

## Discussion

Truncating mutations in *Mybpc3* and RNAi knockdown of cMyBPC levels caused comparable abnormalities in cardiomyocyte contraction and relaxation, supporting the conclusion that *MYBPC3* mutations cause HCM by haploinsufficiency. We show that the severity of cardiomyocyte phenotypes is dependent on cMyBPC levels; truncation of one allele caused milder abnormalities in systolic and diastolic performance than biallelic mutations. The phenotypes of MyBPC-RNAi cardiomyocytes confirmed this dose-dependent relationship; extinguishing post-natal protein expression substantially amplified hypercontractility and impaired relaxation, evidencing a direct role of cMyBPC across the cardiac cycle.

We show that the *MYBPC3* mutations directly alter myosin biophysical properties (Figure 4). Genetic repression of myosin’s motor function, as occurs in Myh6^764/764^ cardiomyocytes,^31^ normalized the hypercontractile phenotype of cMyBPC deficiency, a finding that centrally positions myosin dysregulation as the mechanism for the pathogenicity of cMyBPC mutations. These observations also indicate that strategies to reduce cMyBPC levels can, at least transiently, increase myosin contractility. A second, pharmacological approach to depress myosin function confirmed the conclusion that myosin dysregulation is a unifying mechanism in HCM and indicated the therapeutic potential of targeting myosin in patients with *MYBPC3* mutations as a novel treatment for cMyBPC truncation. As previously shown in HCM mice with myosin mutations^3^, MYK-461- treatment of mouse or human heart tissues with cMyBPC mutations resulted in dose-dependent attenuation of hypercontractility.

**Figure 4:**
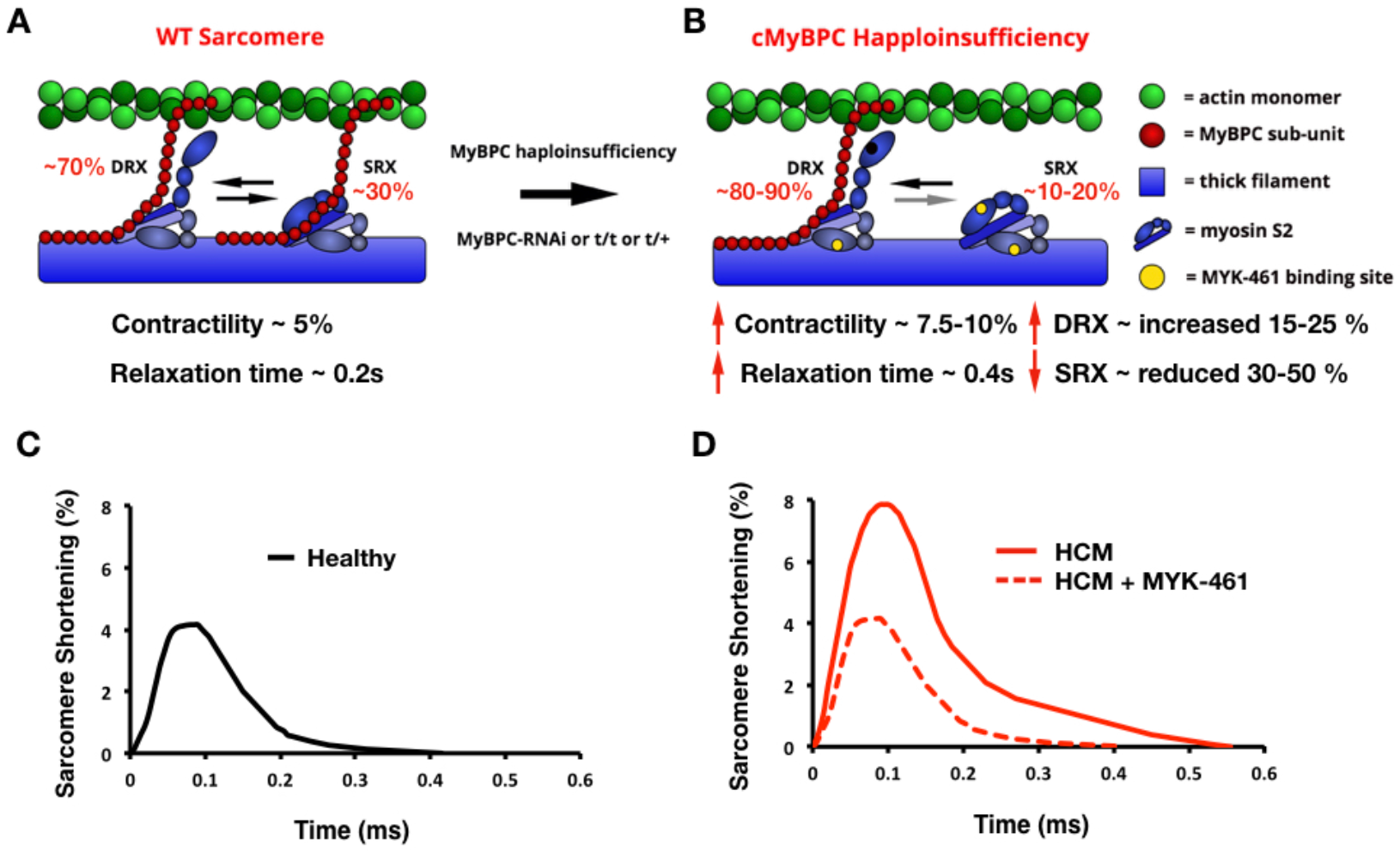
Schematic of the mechanism by which haploinsufficiency of cMyBPC causes HCM. **A)** Schematic of a WT sarcomere with normal cMyBPC levels, and physiologic contractility and relaxation due to appropriate proportions myosins in state of super relaxation (SRX) with low energy consumption or disordered relaxation (DRX) with high energy consumption. **B)** Schematic of an HCM sarcomere with reduced cMyBPC levels that dysregulates the proportions of myosins in DRX (increased) and SRX (reduced). The increased proportion of DRX myosins causes inappropriate sarcomere hypercontractility. Yellow denotes the approximate interaction site of MYK-461 on myosin, which abates the hypercontractile phenotype and shifts the myosin DRX:SRX equilibrium back toward normal. **C)** Contractile waveform of an individual cardiomyocyte isolated from a healthy individual, showing normal sarcomere shortening and normal relaxation duration. **D)** Contractile waveform from a cardiomyocyte isolated from a HCM patient with cMyBPC haploinsufficiency, showing hypercontractility with increased sarcomere shortening and slowed relaxation. MYK-461 normalizes the HCM phenotypes of hypercontractility by restoring physiologic balance of myosin DRX:SRX.

Earlier studies of *Mypbc3*^t/t^ mice using *in vivo* echocardiography or invasive hemodynamic assessments identified reduced systolic contraction, quite unlike the prominent hypercontractility observed in isolated *Mypbc3*^t/t^ or MyBPC-RNAi cardiomyocytes. Several factors may account for these dichotomous findings. *Mypbc3*^t/t^ cardiomyocytes have impaired post-natal cytokinesis that increases numbers of cardiomyocytes, particularly mononuclear cardiomyocytes^25^ that may be dysfunctional. Increased mononuclear cells, hypercontractility of binuclear cells, and altered myocardial geometry may each increase energy demands, which when unmet, could compromise contraction, promote cardiomyocyte death, and increase myocardial fibrosis that is prominent in *Mypbc3*^t/t^ hearts. By eliminating these factors, our analyses of isolated *Mypbc3*^t/t^ and post-natal MyBPC-RNAi cardiomyocytes provided a more proximal readout, revealing that reduced cMyBPC levels increased sarcomere contractility.

cMyBPC deficiency also altered cardiac relaxation. Although we could not demonstrate significant relaxation deficits in *Mypbc3*^t/+^ cardiomyocytes, similar to the normal findings of *Mypbc3*^t/+^ cardiac fibers,^27, 28^ diastolic abnormalities are evident in clinical studies of patients with heterozygous *MYPBC3* mutations^29^ and prominent in *Mypbc3*^t/t^ cardiomyocytes. Both *Myh6*^764/764^ and MYK-461 abrogated increased relaxation durations associated with cMyBPC deficiency, further strengthening the conclusion that myosin dysregulation is the central driver of HCM pathophysiology, in which diastolic dysfunction predominates.

Mant-ATP experiments provided a mechanism for relaxation abnormalities. Our experiments and those by others^27, 28^ show altered proportions of myosins in DRX and SRX in mouse and human myocardium with truncating cMyBPC mutations. Loss of cMyBPC increased the proportions of myosin in the more active DRX conformation. Importantly, MYK-461 restored physiologic DRX/SRX proportions in mice and human tissues, at concentrations that alleviated enhanced cardiomyocyte contractility, data that strongly suggests that increases in the proportion of myosins in DRX contributes to the hyper-contractile phenotype of HCM. As MYK-461 was acutely administered to permeabilized myocardium in our study, its binding to myosin (and not secondary signalling events) is most likely the driver of these beneficial biophysical changes. We therefore suggest that MYK-461 represses filamentous function by directly decreasing myosin contractile function and increasing relaxation properties, providing evidence for dual mechanisms by which MYK-461 can modulate muscle function. These pre-clinical data, reinforced by the data from human myocardium of patients with *Mybpc3* mutations, indicate the potential for MYK-461 to benefit HCM patients with cMyBPC as well as with myosin mutations.^3^

In summation, cMyBPC truncation drives disease by a mechanism of haploinsufficiency, wherein myosin SRX conformations are destabilized, leading to deleterious levels of DRX that drive hyper-contractility, impair relaxation, and increase energy consumption. This triad of abnormalities accurately explains the clinical phenotypes of hyper-dynamic contraction, diastolic dysfunction, and energy deprivation in HCM hearts. By targeting myosin functions, genetically or pharmacologically, these phenotypes are normalized in cardiomyocytes and will likely reduce disease pathogenesis *in vivo*. Demonstrating that myosin is central to the pathophysiology of cMyBPC truncation, and extends the therapeutic utility of MYK-461 to the proportionally largest subset of HCM patients with *MYBPC3* mutations.

## Materials and Methods

### MyBP-C truncation

All animal protocols were compliant with the approved protocols of the Association for the Assessment and Accreditation of Laboratory Animal Care and Harvard Medical School. *Mypbc3*^t/+^, *Mypbc3*^t/t^ and WT mice were studied (129SvEv background) with histopathology being previously described in detail ^23, 29^. The truncated *Mybpc3* alleles were created by PGK-neomycin resistance gene insertion into exon 30 creating a predicted truncation at amino acid 1,064 of the 1,270 residues of cMyBPC. The homozygous truncated allele has been shown to produce less than 10% of normal cMyBPC. It should be noted that a greater proportion of cells in the *Mypbc3*^t/t^ cohort exhibited fibrillation and were excluded from contractile measures as they could not be reliably paced, making measures in *Mypbc3*^t/t^ cardiomyocytes more challenging.

### RNAi

RNAi was delivered at P10 by AAV vector using AAV9 capsid packaging by triple transfection as described^25^. The AAV9 vector contains an shRNA construct that specifically targets specific 21-base-pair sequence targeted to *Mybpc3* exon19 and an enhanced GFP plasmid (Addgene). Vector was injected into the thoracic cavity of P1 t/+ neonates [5 × 10^13^ viral genomes (vg)/kg], GFP fluorescence was used to identify isolated cardiomyocytes that had taken up the vector. GFP fluorescence was evident from 48 hours post-injection for at least 5 months, RNAi reduced cMyBPC expression to ∼10% of normal ^25^.

### Human myectomy samples

Human myectomy samples were obtained after written informed consent from three HCM patients with distinct heterozygous frameshift truncating variants (Gln981fs, Leu1014fs, and Lys1209fs) in *MYBPC3*. Myectomy samples of the septum were flash frozen and stored in liquid nitrogen and tissue preparation for Mant-ATP experiments were performed as described below.

### Myocyte isolation

Cardiomyocytes were isolated from 8-20 week old mice by rapid explantation and aortic cannulation on a Langendorff apparatus for perfusion with Enzyme Buffer (EB composition: 135 mM NaCl, 4 mM KCl, 0.33 mM NaH_2_PO_4_, 1 mM MgCl_2_, and 10 mM HEPES, pH 7.40, which incorporated Collagenase D, Collagenase B and Protease XIV) for 10 minutes. After perfusion the atria and right ventricle were removed and the left ventricle was minced in TA buffer (Composition: 135 mM NaCl, 4 mM KCl, 0.33 mM NaH_2_PO_4_, 1 mM MgCl_2_, and 10 mM HEPES, pH 7.40, which incorporated bovine serum albumin) and passed through a 100 μm filter into a 50ml conical tube. Tissue settled for 15 minutes to allow myocytes to pellet by gravity. The pellet was then sequentially resuspended every 10 minutes through an increasing calcium gradient (5%, 20%, 50%, 100% calcium tyrode) to provide a cell fraction enriched in myocytes with a final experimental buffer (Composition: NaCl 137 mM, KCl 5.4 mM, CaCl_2_ 1.2 mM, MgCl_2_ 0.5 mM, HEPES 10 mM at pH 7.4, which incorporated glucose).

### Contractile measures of myocyte function

Isolated cardiomyocytes were placed in wells of a 6-well plate that had been precoated with laminin. Laminin coating was performed for two hours before cardiomyocyte introduction at a concentration of 10 ug/mL in PBS (Composition: KH_2_PO_4_ 1mM, NaCl 155 mM, Na_2_HPO_4_ 3mM, at pH 7.4). Laminin coating solution was washed once with PBS before cells were introduced into the wells. Once cells were introduced they were left to incubate for 10 minutes to equilibrate to experimental temperature (30°C). Cells were imaged using a Keyence BZ-X710 microscope using a Nikon 40X/0.65 NA objective. Cells were kept at 30°C using microscope specific incubation chamber that was also used to deliver 20% O_2_ and 5% CO_2_ to the experimental chamber. Cells were paced at 1Hz using custom-built electrodes hooked up to a pacing unit (Pulsar 6i, FHC Brunswick, ME, USA) delivering 20V. Movies were acquired at 29 frames per second for 5 seconds (5 contractile cycles).

An ImageJ plugin SarCoptiM was used to track sarcomere lengths during contractile cycles ^35^. Sarcomere tracking was then used to calculate cellular shortening (%), relaxed and contracted sarcomere lengths (μm), contractile cycle and relaxation durations (seconds).

For experiments incorporating MYK-461, drug was applied in concentrations ranging 0. 05 – 0.3 μM in the experimental buffer. MYK-461 was incubated with cells for a minimum of 10 minutes before data acquisition.

### Mant-ATP experiments

Mice were sacrificed by rapid cervical dislocation, atria and right ventricle were removed and samples were flash frozen in liquid nitrogen. Mant-ATP protocols were adapted from publications^27,34^: 20 mg left ventricular human or mouse myocardial samples were thawed in permeabilization buffer (Composition: NaCl 100 mM, MgCl_2_ 8 mM, EGTA 5 mM, K_2_HPO_4_ 5 mM, KH_2_PO_4_ 5 mM, NaN_3_ 3 mM, ATP 5 mM, DTT 1 mM, BDM 20 mM, Triton-X 100 0.1%, at pH 7.0). Samples were permeabilized for 6 hours on ice on a rocker solution changes occurring every two hours. At the completion of this step samples were stored overnight at −20°C in glycerinating solution (Composition: K acetate, 120 mM; Mg acetate, 5 mM; K_2_HPO_4_, 2.5 mM; KH_2_PO_4_, 2.5 mM; MOPS, 50 mM; ATP, 5 mM; BDM, 20 mM; DTT, 2 mM; glycerol, 50% (v/v), pH 6.8.) for dissection within 2 days.

Once glycerinated ventricular myocardium was dissected into ∼ 90 × 400 μm pieces that were held under two pins in a chamber constructed from a slide and coverslip. These samples were permeabilized using the permeabilization buffer for a further 30 minutes on ice prior to experimentation. After secondary permeabilization chambers were flushed with glycerinating buffer.

For fluorescence acquisition a Nikon TE2000-E was used with a Nikon 20X/0.45 objective, using a Hammamatsu C9100 EM-CCD. Frames were acquired every 10 seconds with a 20 ms acquisition and exposure time using a DAPI filter set, images were collected for 15 minutes. Prior to acquisition each chamber was flushed with ATP buffer (Composition: K acetate 120 mM, Mg acetate 5 mM, K_2_HPO_4_ 2.5 mM, KH_2_PO_4_ 2.5 mM, ATP 4 mM, MOPS 50 mM, DTT 2 mM at pH 6.8) to remove glycerol. This buffer was replaced with two chamber volumes of rigor buffer (Composition: K acetate 120 mM, Mg acetate 5 mM, K_2_HPO_4_ 2.5 mM, KH_2_PO_4_ 2.5 mM, MOPS 50 mM, DTT 2 mM at pH 6.8). Rigor buffer was incubated for 5 minutes to allow rigor to set in. Initial fluorescence acquisition was simultaneous with the addition of one chamber volume of rigor buffer with 250 μM Mant-ATP to visualize fluorescent Mant-ATP wash in. At the end of a 15-minute acquisition, a chamber volume of ATP buffer (Rigor buffer + 4 mM ATP) was added to the chamber with simultaneous acquisition of the Mant-ATP chase. For experiments with MYK-461 all experimental solutions contained 0.3 μM MYK-461.

### Mant-ATP analysis

Similar to protocols previously described for analysis^27^ three regions of each myocardial tissue strip were sampled for fluorescence decay using the ROI manager in ImageJ. The final data point of fluorescence wash in defined the y-intercept. All data was plot as a normalized intensity of initial fluorescent intensity from the three sampled regions. These data are fit to an unconstrained double exponential decay using Sigmaplot:

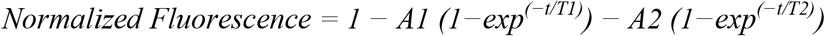

Where A1 is the amplitude of the initial rapid decay approximating the ‘disordered relaxed state’ (DRX) with T1 as the time constant for this decay. A2 is the slower second decay approximating the proportion of myosin heads in the ‘super relaxed state’ (SRX) with its associated time constant T2.

Each individual experiment was fit using this double exponential decay with all values determined and plot. Statistical analysis was performed using 2-way ANOVA with multiple comparisons tests.

### Statistics

Where appropriate students t-tests were employed. In instances where experimental hypothesis were tested amongst multiple treatment groups one-way analysis of variance (ANOVA) was used. For multiple comparisons post-hoc Bonferroni corrections were used with a significance cut off of p < 0.05.

## Supplementary Materials

1. Supplementary Materials and Methods
  Quantification of cMyBPC protein levels in *Mybpc3* mouse models
  *In vivo* comparisons of *Mybpc3* mouse models
  Characterization of viral titers and depletion of *Mybpc3* transcripts
  Assessing fluorescent decay in Mant-ATP experiments
  Mant-ATP control measures to control for hypertrophy or myosin DCM on assay findings
2. Supplementary Figures
  Fig. S1. Representation of *Mybpc3* mouse models
  Fig. S2. *In vivo* cardiac function and proteomic characterization in *Mybpc3* mouse models
  Fig. S3. The protein and function effects of increasing MyBPC-RNAi titers
  Fig. S4. Analysis of Mant-ATP video files
  Fig. S5. Mant-ATP assays in cardiac tissues for *Myh6*^764/764^ mice with DCM
3. Supplementary Movie
  S1. Mant-ATP fluorescence decay

## Acknowledgments

We thank Dr. Roger Cooke for assistance with the Mant-ATP assays. This work was supported in part by grants from a Sir Henry Wellcome postdoctoral fellowship from the Wellcome Trust (206466/Z/17/Z C.N.T.) the Sarnoff Foundation (A.G.), National Medical Research Council (NMRC) and Ministry of Education, Singapore for D.L. and J.J., American Heart Association Midwest Predoctoral Fellowship (15PRE22430028 to TLL), National Institutes of Health grants (R01HL130356, R01HL105826, and K02HL114749 to SS),Research council of Norway (221707 to IGL). British Heart Foundation (Programme grant RG/12/16/29939) and the British Heart Foundation Centre of Research Excellence (Oxford) to H.W. and C.S.R. The National Heart Blood and Lung Institutes (HL084553 and HL080494 to J.G.S. and C.E.S.), and the Howard Hughes Medical Institute (C.E.S.)

## Author contributions

C.N.T. and H.W. performed experiments on mouse cardiomyocytes, H.W. performed in-vivo characterization of mice and RNAi injection. D.L. and J.J. produced the RNAi virus. Quantification of Mybpc3 expression was performed by A.T. and J.G. Mouse genotyping was performed by H.W. and M.L. Preparation of Human myectomy samples was performed by B.M. and C.N.T. C.N.T., A.C.G., and H.W. performed with Mant-ATP experiments. Blots of cMyBPC were performed by H.W., T.L.L and S.S. Intellectual design of experiments C.N.T., H.W., J.J., I.G.L., C.E.S., J.G.S. Editing and preparation of manuscript all authors.

## References and Notes

1. Maron BJ, Gardin JM, Flack JM, Gidding SS, Kurosaki TT and Bild DE. Prevalence of hypertrophic cardiomyopathy in a general population of young adults. Echocardiographic analysis of 4111 subjects in the CARDIA Study. Coronary Artery Risk Development in (Young) Adults. Circulation. 1995; 92:785–9.

2. Maron BJ, Casey SA, Poliac LC, Gohman TE, Almquist AK and Aeppli DM. Clinical course of hypertrophic cardiomyopathy in a regional United States cohort. JAMA. 1999; 281:650–5.

3. Green EM, Wakimoto H, Anderson RL, Evanchik MJ, Gorham JM, Harrison BC, Henze M, Kawas R, Oslob JD, Rodriguez HM, Song Y, Wan W, Leinwand LA, Spudich JA, McDowell RS, Seidman JG and Seidman CE. A small-molecule inhibitor of sarcomere contractility suppresses hypertrophic cardiomyopathy in mice. Science. 2016; 351:617–21.

4. Walsh R, Thomson KL, Ware JS, Funke BH, Woodley J, McGuire KJ, Mazzarotto F, Blair E, Seller A, Taylor JC, Minikel EV, Exome Aggregation C, MacArthur DG, Farrall M, Cook SA and Watkins H. Reassessment of Mendelian gene pathogenicity using 7,855 cardiomyopathy cases and 60,706 reference samples. Genet Med. 2017; 19:192–203.

5. Moolman-Smook JC, De Lange WJ, Bruwer EC, Brink PA and Corfield VA. The origins of hypertrophic cardiomyopathy-causing mutations in two South African subpopulations: a unique profile of both independent and founder events. Am J Hum Genet. 1999; 65:1308–20.

6. Kubo T, Kitaoka H, Okawa M, Matsumura Y, Hitomi N, Yamasaki N, Furuno T, Takata J, Nishinaga M, Kimura A and Doi YL. Lifelong left ventricular remodeling of hypertrophic cardiomyopathy caused by a founder frameshift deletion mutation in the cardiac Myosin-binding protein C gene among Japanese. J Am Coll Cardiol. 2005; 46:1737–43.

7. Teirlinck CH, Senni F, Malti RE, Majoor-Krakauer D, Fellmann F, Millat G, Andre-Fouet X, Pernot F, Stumpf M, Boutarin J and Bouvagnet P. A human MYBPC3 mutation appearing about 10 centuries ago results in a hypertrophic cardiomyopathy with delayed onset, moderate evolution but with a risk of sudden death. BMC Med Genet. 2012; 13:105.

8. Saltzman AJ, Mancini-DiNardo D, Li C, Chung WK, Ho CY, Hurst S, Wynn J, Care M, Hamilton RM, Seidman GW, Gorham J, McDonough B, Sparks E, Seidman JG, Seidman CE and Rehm HL. Short communication: the cardiac myosin binding protein C Arg502Trp mutation: a common cause of hypertrophic cardiomyopathy. Circ Res. 2010; 106:1549–52.

9. Sabater-Molina M, Saura D, Garcia-Molina Saez E, Gonzalez-Carrillo J, Polo L, Perez-Sanchez I, Olmo MD, Oliva-Sandoval MJ, Barriales-Villa R, Carbonell P, Pascual-Figal D and Gimeno JR. A Novel Founder Mutation in MYBPC3: Phenotypic Comparison With the Most Prevalent MYBPC3 Mutation in Spain. Rev Esp Cardiol (Engl Ed). 2017; 70:105–114.

10. Adalsteinsdottir B, Teekakirikul P, Maron BJ, Burke MA, Gudbjartsson DF, Holm H, Stefansson K, DePalma SR, Mazaika E, McDonough B, Danielsen R, Seidman JG, Seidman CE and Gunnarsson GT. Nationwide study on hypertrophic cardiomyopathy in Iceland: evidence of a MYBPC3 founder mutation. Circulation. 2014; 130:1158–67.

11. van Velzen HG, Schinkel AFL, Oldenburg RA, van Slegtenhorst MA, Frohn-Mulder IME, van der Velden J and Michels M. Clinical Characteristics and Long-Term Outcome of Hypertrophic Cardiomyopathy in Individuals With a MYBPC3 (Myosin-Binding Protein C) Founder Mutation. Circ Cardiovasc Genet. 2017; 10.

12. Dhandapany PS, Sadayappan S, Xue Y, Powell GT, Rani DS, Nallari P, Rai TS, Khullar M, Soares P, Bahl A, Tharkan JM, Vaideeswar P, Rathinavel A, Narasimhan C, Ayapati DR, Ayub Q, Mehdi SQ, Oppenheimer S, Richards MB, Price AL, Patterson N, Reich D, Singh L, Tyler-Smith C and Thangaraj K. A common MYBPC3 (cardiac myosin binding protein C) variant associated with cardiomyopathies in South Asia. Nat Genet. 2009; 41:187–91.

13. Marston S, Copeland O, Jacques A, Livesey K, Tsang V, McKenna WJ, Jalilzadeh S, Carballo S, Redwood C and Watkins H. Evidence from human myectomy samples that MYBPC3 mutations cause hypertrophic cardiomyopathy through haploinsufficiency. Circ Res. 2009; 105:219–22.

14. van Dijk SJ, Dooijes D, dos Remedios C, Michels M, Lamers JM, Winegrad S, Schlossarek S, Carrier L, ten Cate FJ, Stienen GJ and van der Velden J. Cardiac myosin-binding protein C mutations and hypertrophic cardiomyopathy: haploinsufficiency, deranged phosphorylation, and cardiomyocyte dysfunction. Circulation. 2009; 119:1473–83.

15. Ho CY, Sweitzer NK, McDonough B, Maron BJ, Casey SA, Seidman JG, Seidman CE and Solomon SD. Assessment of diastolic function with Doppler tissue imaging to predict genotype in preclinical hypertrophic cardiomyopathy. Circulation. 2002; 105:2992–7.

16. Forsey J, Benson L, Rozenblyum E, Friedberg MK and Mertens L. Early changes in apical rotation in genotype positive children with hypertrophic cardiomyopathy mutations without hypertrophic changes on two-dimensional imaging. J Am Soc Echocardiogr. 2014; 27:215–21.

17. Russel IK, Brouwer WP, Germans T, Knaapen P, Marcus JT, van der Velden J, Gotte MJ and van Rossum AC. Increased left ventricular torsion in hypertrophic cardiomyopathy mutation carriers with normal wall thickness. J Cardiovasc Magn Reson. 2011; 13:3.

18. Viswanathan SK, Sanders HK, McNamara JW, Jagadeesan A, Jahangir A, Tajik AJ and Sadayappan S. Hypertrophic cardiomyopathy clinical phenotype is independent of gene mutation and mutation dosage. PLoS One. 2017; 12:e0187948.

19. Trivedi DV, Adhikari AS, Sarkar SS, Ruppel KM and Spudich JA. Hypertrophic cardiomyopathy and the myosin mesa: viewing an old disease in a new light. Biophys Rev. 2017.

20. Alamo L, Ware JS, Pinto A, Gillilan RE, Seidman JG, Seidman CE and Padron R. Effects of myosin variants on interacting-heads motif explain distinct hypertrophic and dilated cardiomyopathy phenotypes. Elife. 2017; 6.

21. Pfuhl M and Gautel M. Structure, interactions and function of the N-terminus of cardiac myosin binding protein C (MyBP-C): who does what, with what, and to whom? J Muscle Res Cell Motil. 2012; 33:83–94.

22. Kensler RW, Shaffer JF and Harris SP. Binding of the N-terminal fragment C0-C2 of cardiac MyBP-C to cardiac F-actin. J Struct Biol. 2011; 174:44–51.

23. McConnell BK, Fatkin D, Semsarian C, Jones KA, Georgakopoulos D, Maguire CT, Healey MJ, Mudd JO, Moskowitz IP, Conner DA, Giewat M, Wakimoto H, Berul CI, Schoen FJ, Kass DA, Seidman CE and Seidman JG. Comparison of two murine models of familial hypertrophic cardiomyopathy. Circ Res. 2001; 88:383–9.

24. Previs MJ, Mun JY, Michalek AJ, Previs SB, Gulick J, Robbins J, Warshaw DM and Craig R. Phosphorylation and calcium antagonistically tune myosin-binding protein C’s structure and function. Proc Natl Acad Sci U S A. 2016; 113:3239–44.

25. Jiang J, Burgon PG, Wakimoto H, Onoue K, Gorham JM, O’Meara CC, Fomovsky G, McConnell BK, Lee RT, Seidman JG and Seidman CE. Cardiac myosin binding protein C regulates postnatal myocyte cytokinesis. Proc Natl Acad Sci U S A. 2015; 112:9046–51.

26. Korte FS, McDonald KS, Harris SP and Moss RL. Loaded shortening, power output, and rate of force redevelopment are increased with knockout of cardiac myosin binding protein-C. Circ Res. 2003; 93:752–8.

27. McNamara JW, Li A, Smith NJ, Lal S, Graham RM, Kooiker KB, van Dijk SJ, Remedios CGD, Harris SP and Cooke R. Ablation of cardiac myosin binding protein-C disrupts the superrelaxed state of myosin in murine cardiomyocytes. J Mol Cell Cardiol. 2016; 94:65–71.

28. McNamara JW, Li A, Lal S, Bos JM, Harris SP, van der Velden J, Ackerman MJ, Cooke R and Dos Remedios CG. MYBPC3 mutations are associated with a reduced super-relaxed state in patients with hypertrophic cardiomyopathy. PLoS One. 2017; 12:e0180064.

29. McConnell BK, Jones KA, Fatkin D, Arroyo LH, Lee RT, Aristizabal O, Turnbull DH, Georgakopoulos D, Kass D, Bond M, Niimura H, Schoen FJ, Conner D, Fischman DA, Seidman CE and Seidman JG. Dilated cardiomyopathy in homozygous myosin-binding protein-C mutant mice. J Clin Invest. 1999; 104:1235–44.

30. Watkins H, Conner D, Thierfelder L, Jarcho JA, MacRae C, McKenna WJ, Maron BJ, Seidman JG and Seidman CE. Mutations in the cardiac myosin binding protein-C gene on chromosome 11 cause familial hypertrophic cardiomyopathy. Nat Genet. 1995; 11:434–7.

31. Schmitt JP, Debold EP, Ahmad F, Armstrong A, Frederico A, Conner DA, Mende U, Lohse MJ, Warshaw D, Seidman CE and Seidman JG. Cardiac myosin missense mutations cause dilated cardiomyopathy in mouse models and depress molecular motor function. Proc Natl Acad Sci U S A. 2006; 103:14525–30.

32. Kamisago M, Sharma SD, DePalma SR, Solomon S, Sharma P, McDonough B, Smoot L, Mullen MP, Woolf PK, Wigle ED, Seidman JG and Seidman CE. Mutations in sarcomere protein genes as a cause of dilated cardiomyopathy. N Engl J Med. 2000; 343:1688–96.

33. Barefield D, Kumar M, Gorham J, Seidman JG, Seidman CE, de Tombe PP and Sadayappan S. Haploinsufficiency of MYBPC3 exacerbates the development of hypertrophic cardiomyopathy in heterozygous mice. J Mol Cell Cardiol. 2015; 79:234–43.

34. Hooijman P, Stewart MA and Cooke R. A new state of cardiac myosin with very slow ATP turnover: a potential cardioprotective mechanism in the heart. Biophys J. 2011; 100:1969–76.

35. Pasqualin C, Gannier F, Yu A, Malecot CO, Bredeloux P and Maupoil V. SarcOptiM for ImageJ: high-frequency online sarcomere length computing on stimulated cardiomyocytes. Am J Physiol Cell Physiol. 2016; 311:C277–83.

